# DLPFC-PPC-cTBS Effects on Metacognitive Awareness

**DOI:** 10.1101/2022.10.26.513949

**Authors:** Antonio Martin, Timothy J. Lane, Tzu-Yu Hsu

## Abstract

**Background:** Neuroimaging and lesion studies suggested that the dorsolateral prefrontal and posterior parietal cortices mediate visual metacognitive awareness. The causal evidence provided by non-invasive brain stimulation, however, is inconsistent.

**Objective/Hypothesis:** Here we revisit a major figure discrimination experiment adding a new Kanizsa figure task trying to resolve whether bilateral continuous theta-burst transcranial magnetic stimulation (cTBS) over these regions affects perceptual metacognition. Specifically, we tested whether subjective visibility ratings and/or metacognitive efficiency are lower when cTBS is applied to these two regions in comparison to an active control region.

**Methods:** A within-subjects design including three sessions spaced by one-week intervals was implemented. In each session, every participant was administered bilateral cTBS to either prefrontal, control or parietal cortices. Two concurrent tasks were performed, a real and an illusory figure task, stabilising objective performance with use of an adaptive staircase procedure.

**Results:** When performing the replicated task, cTBS was found insufficient to disrupt neither visibility ratings nor metacognitive efficiency. However, with use of Kanizsa style illusory figures, cTBS over the dorsolateral prefrontal, but not over the posterior parietal cortex, was observed to significantly diminish metacognitive efficiency.

**Conclusion(s):** Real and illusory figure tasks demonstrated different cTBS effects. A possible explanation is the involvement of the prefrontal cortex in the creation of expectations, which is necessary for efficient metacognition. Failure to replicate previous findings for the real figure task, however, cannot be said to support, conclusively, the notion that these brain regions have a causal role in metacognitive awareness. This inconsistent finding may result from certain limitations of our study, thereby suggesting the need for yet further investigation.

## Introduction

Visual awareness of perceptual contents has been suggested to require the involvement of first person monitoring of one’s own mental processing (Brown et al., 2019). One approach to evaluating visual awareness is use of metacognitive judgements (i.e., subjective ratings), including visibility or confidence, along with objective performance. Such an approach can be implemented in various cognitive experiments, including visual discrimination tasks, which allow us to investigate metacognitive sensitivity, or how well metacognitive judgements can predict objective performance. Several brain regions in the prefrontal parietal network (PPN; Bor & Seth, 2012) have shown to be involved in metacognitive judgements in various perception (Cai et al., 2022; Fleming et al., 2010, 2012, 2015; Lapate et al., 2020; Miyamoto et al., 2021; Rahnev et al., 2016; Shekhar and Rahnev, 2018) or memory based cognitive tasks (Cai et al., 2022; Ryals et al., 2016; Yazar et al., 2014; Ye et al., 2018, 2019). Several PPN brain regions overlap with the central executive network (Ryali et al., 2016) or the lateral frontoparietal network (Uddin et al., 2019); in each case the relevant regions include the dorsolateral prefrontal and the posterior parietal cortices (DLPFC and PPC respectively). However, causal studies--as implemented by non-invasive brain stimulation (NIBS)--which suggest that the DLPFC mediates metacognitive judgements (Shekhar & Rahnev, 2018) or metacognitive sensitivity, (Rounis et al., 2010; Ruby et al., 2018) have been challenged (Bor et al., 2017, 2018). Ruby et al. (2018) addressed these challenges both conceptually and via simulations, pointing out that low statistical power can produce false negatives, even when a true positive effect has been identified. To the best of our knowledge, this challenge has not been addressed adequately yet; optimally, further enhanced empirical investigation is required, taking into account improved statistical power, both through additional participants and improved experimental design.”

A first line of evidence derives from correlational and lesion studies that show patients with a damaged prefrontal cortex who have intact objective performance but selective impairment of metacognitive awareness (Colás et al., 2019; Del Cul et al., 2009; Grossner et al., 2018). For example, Colás et al. (2019) observed that patients with DLPFC damage, despite having normal levels of objective spatial orienting abilities, presented a more conservative response bias affecting their subjective visibility threshold. Furthermore, parietal cortex neural activity, especially the PPC, is also associated with confidence level (Hanks et al., 2011; Kiani & Shadlen, 2009; Pereira et al., 2020; Simons et al., 2010). Simons et al. (2010) observed specific decreases in confidence levels after bilateral impairments to the PPC, similar to those described for the DLPFC damages. Although these results suggest the PPN as essential to metacognitive tasks, the linkage of the PPC to processes parallel to those of the DLPFC still does not show clear evidence on whether they both correlate redundantly to overall response bias or metacognitive sensitivities.

A second line of evidence derived from NIBS studies further supports the causal role played by the DLPFC in metacognitive awareness. Rounis and her colleagues (2010) applied bilateral offline continuous theta-burst transcranial magnetic stimulation (cTBS) to temporarily disrupt the PPN activity, specifically in the DLPFC, in order to evaluate DLPFC’s influence on metacognitive sensitivity. They observed that bilateral DLPFC cTBS affected visibility ratings, for accurate objective performance: that is to say that although their performance was accurate, their visibility ratings were impaired. This dissociation between performance and visibility is an instance of impaired metacognitive sensitivity. The authors hypothesised that the DLPFC is causally involved in metacognitive sensitivity. An opposing view derives from Shekhar and Rahnev (2018), who used online transcranial magnetic stimulation (TMS) and concluded that the anterior PFC--but not the DLPFC--plays a central role in metacognitive sensitivity. They did find, however, that the DLPFC modulates all confidence ratings. Several studies also showed supporting and consistent findings (Fleming et al., 2010, 2012, 2014; Ryals et al., 2016; Rahnev et al., 2016). Based upon this finding they hypothesised that the DLPFC causally affects the strength of the sensory evidence. Strikingly, Bor et al. (2017), in both between- and within-subjects failed to replicate any of these findings, suggesting that neither the DLPFC nor the PPC are playing the hypothesised roles.

Several methodological differences were pointed out as potentially explaining the failure in replicating the effects of cTBS in metacognition, particularly the exclusion of unstable datasets (Ruby et al., 2018; Bor et al., 2018). In this piece of work, we aimed to refine the experimental methodology and raise questions on whether the task originally used is suitable for investigating the effects of cTBS over metacognitive awareness. For this aim we decided to compare the original task to a similar yet new task using Kanizsa illusory contours to potentially elevate the effects of cTBS over metacognition. The DLPFC has been suggested to contribute to the formation of expectations that shape subsequent responses, particularly encoding the reliability of expected sensory inputs (de Lange et al., 2018; Lobanov et al., 2014; Tanaka et al., 2006). According to the predictive coding hypothesis (Friston & Kiebel, 2009), when bottom-up sensory inputs are ambiguous or unreliable --as with illusory contours--there is a tendency to give more weight to information from internal models (Parr et al., 2018; Yu, 2014). Under this hypothesis, altering the DLPFC by means of cTBS should highly disrupt metacognition when stimuli are ambiguous (Sherman et al., 2015) or highly dependent on internal models, such as when perceiving illusory contours, in comparison to more readily available stimuli, such as when using real figures

In sum, the question of whether the DLPFC or the PPC cTBS negatively affects metacognitive awareness remains to be resolved. In the present study, we examine how the PPN is related to metacognitive visual awareness. In order to do so, we replicated the Rounis et al. (2010) figure discrimination task and compared it to an illusory Kanizsa figure task (Kanizsa, 1976). When three or four ‘pacmen’-like inducers are arranged in a configuration, they can create the illusion that a shape exists, even without an actual physical shape. We chose Kanizsa style figures because of their similarity to the original task and because they are assumed to depend on top-down contextual neural activities associated with unreliable bottom-up sensory inputs (Banica & Schwarzkopf, 2016; Kok et al., 2016). Adhering to a within-subjects design, offline cTBS was administered bilaterally at either the DLPFC or the PPC, during different sessions, and then compared to an active control site, the somatosensory cortex (S1).

## Method

### Participants

All data exclusions and inclusion criteria were established prior to data analysis, all manipulations, and all measures in the study (https://osf.io/78b3x/). The only one we cannot meet was the sample size due to the COVID19 outbreak and limitation of funding. Although a total of thirty-five participants were recruited for this study after meeting our inclusion criteria (see Supplementary information S2), two of them could not finish the experiment due to a COVID19 outbreak. Although thirty-three participants successfully completed the experiment, twelve met our exclusion criteria by presenting low task performance or unstable datasets (see Supplementary information S3). Our final list was constituted by 11 females and 10 males, with ages ranging from 23 to 41 with an average of 30.92 (SD = 5.23). Age was not statistically different between genders (t_17.61_ = 1.41, p = 0.18). All participants had normal or corrected-to-normal vision. Written consent was to be obtained from all participants prior to their participation in this study. All participants were informed about the purpose of the study and procedures before being asked to give consent. This study was approved by the joint institutional Review Board, Taipei Medical University, Taipei City, Taiwan (N201910050). Anonymised processed data are available at https://osf.io/78b3x. The conditions of our ethics approval do not permit public archiving of raw data. To access this raw data please contact the corresponding author. Access will be granted to named individuals in accordance with ethical procedures governing the reuse of sensitive data. Specifically, requestors must undergo the completion of an agreement on the formal data sharing which is approved by the local IRB.

### Apparatus

Stimuli were generated and delivered with PsychoPy2 Experiment Coder v1.90.3 (Peirce et al., 2019), running under Python on an LCD monitor with a vertical refresh rate of 60 Hz. Participants viewed the stimuli with their head position stabilised by a chin-rest, at a distance of 40 cm. All statistical analyses were performed with R 4.0.3 and JASP 0.16.3. All the experimental stimuli and presentation script was pre-registered prior the research being conducted and available on the open science framework (OSF) repository (https://osf.io/78b3x/).

### Stimuli and visual discrimination tasks

The Rounis et al. (2010) experimental design was adopted with minor adjustments following Bor et al. (2017) design described at the Procedure section. Participants were required to perform a two-alternative forced choice based visual discrimination task (see Figure 1A) in a dark room. In the real figure task, two 0.8° visual angle black stimuli, a diamond and a square, were presented simultaneously, on a grey background, for 33 ms on the left and right respectively, 1^0^ visual angle away from the central fixation cross and followed by a mask for 50 ms. Participants were asked to discriminate the location of a square or a diamond, either on the left or right, while fixating their gaze on a fixation cross in the centre of the screen. In the illusory figure task, the stimuli were replaced by 0.425° radius wide ‘pacmen’-like inducers at every vertex to induce 0.8° wide square-like or diamond-like Kanizsa illusory shapes for 83 ms (see Figure 1B), also followed by a mask for 50 ms. Giving similar instructions at both tasks, participants were asked to discriminate whether the square was on the left and the diamond on the right (left keys: ‘c’ or ‘v’), or vice versa (right keys: ‘n’ or ‘m’), while fixating their gaze on the fixation cross in the centre of the screen. The location of these opposing figures was pseudo randomised with equal probability. Simultaneously, participants were asked to judge the visibility of the stimuli presented relative to how it typically looks up to that moment. If the stimuli at the current trial were sensed as more readily visible compared to other stimuli seen in the experimental context, the participant was asked to report “clear” by pressing keys ‘c’ (if the target presented on the left visual field) or ‘m’ (if the target presented on the right visual field). Alternatively, they were asked to press the “unclear” button ‘v’ (if the target presented on the left visual field) or ‘n’ (if the target presented on the right visual field).

**Figure 1.**
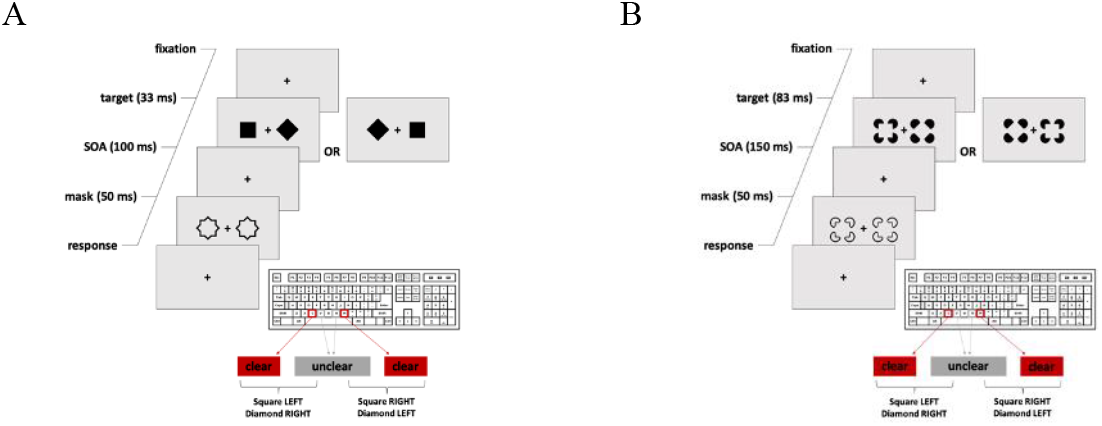
Experimental Design. (A) Replicated two-alternative forced-choice visual task (from Rounis et al., 2010). Participants were asked to discriminate the location of the square to be in either left or right to the fixation cross. Simultaneously, participants also offer a rating over how strong is their subjective visibility (“clear” or “unclear”). (B) New version of the visual task. In this second experiment, participants were given the same instructions but need to locate the illusory square.

We decided to keep with the original instructions agreeing with King & Dehaene (2014) in that visibility ratings might not be exactly identical to confidence ratings when one desires to study conscious perception. Although several parameters have been developed to assess metacognitive sensitivity (Fleming & Lau, 2014), we followed the meta-d’ parameter estimation since it has been argued to be invariant to response biases and to provide a more transparent measure of metacognitive ability (Maniscalco & Lau, 2012). In order to calculate the meta-d’ values, the visibility reports were taken to generate approximations of what the type1 d’ values would be with use of the sum of squared errors approximation (SSE method).

### Theta-burst stimulation

The cTBS pulses were administered with a Magpro X100 (MagVenture, Denmark) in a 70 mm figure-of-eight-shaped coil (MC-B70, MagVenture). Each bilateral cTBS session comprises 600 pulses, at 80% intensity stimulation for adjusted active motor threshold (adjAMT), and were applied over the left and right DLPFC, PPC, or S1.

Every experimental session comprises the administration of cTBS to one region bilaterally. The sequence of stimulation was counterbalanced for each participant across the three experimental sessions. The MNI coordinates for bilateral DLPFC were right [36 30 50] and left [-42 20 50] (Cieslik et al., 2013; Scheperjans et al., 2008); for bilateral PPC were right [36 -53 50] and left [-34 -52 50]; and for the S1 (control site) were right [22 -32 70] and left [-22 -30 70] (see Figure 2). Anatomically, bilateral DLPFC corresponded to the range of coordinates described in Rounis et al (2010), Rahnev et al (2016), and Shekhar & Rahnev (2018). An average observed adjAMT of 28.92 (SD = 5.81) led to cTBS intensities of 24.44 (SD = 7.32) for the DLPFC, 41.96 (SD = 9.07) for the PPC and 47.83 (SD = 9.91) for the S1. More information regarding how the adjAMT was calculated for each region depending on its depth can be found in Supplementary information (S1).

**Figure 2.**
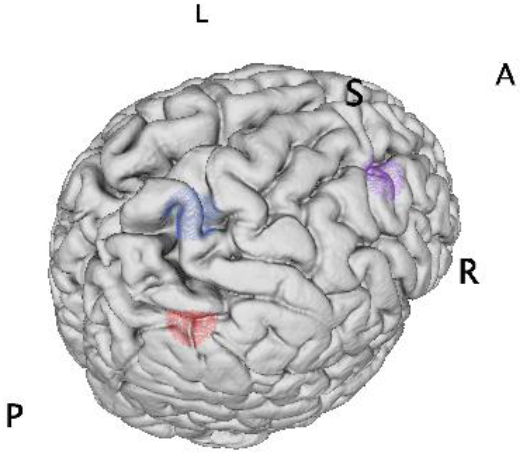
cTBS Stimulation Sites. Right hemisphere view of bilaterally targeted sites: DLPFC (purple), PPC (red), and S1 (blue).

cTBS delivers a 20 seconds train of uninterrupted theta-bursts pulses. This consists of 3 pulses, at 50 Hz, given in 200 ms intervals, comprising a total of 300 pulses for 20 seconds (or 100 bursts), which was applied to one hemisphere. After one-minute inter-stimulation interval, the procedure was repeated in the corresponding location of the other hemisphere. The cTBS protocol followed Rounis et al (2010) and Bor et al (2017). A 300-pulse, 20-second thcTBS protocol has been shown to suppress activity in the stimulated brain region for up to 20 minutes following stimulation (Huang et al., 2005). The coil was attached perpendicular to the location of stimulation (90^0^ angle) and the task would be initiated about 1 minute after cTBS administration.

### Magnetic resonance imaging

Before participants begin the three main experimental sessions, anatomical images for each of them were collected, using a 3T magnetic resonance scanner either Siemens MAGNETOM Skyra 3T or GE 3.0 Tesla Discovery MR750 depending on accessibility. Their T1-weighted images were used to locate the regions for administration of the cTBS as well as for adjAMT calculations.

### Procedure

This study was carried out over four days, with one-week intervals. On the first day, participants were introduced to the rationale of the experiment and did practice trials to ensure their understanding of this study. After providing their consent to participate, both their anatomical images and their AMT were collected. In the following three days, participants were required to do as many practice blocks as necessary, consisting of 100 trials each, in order to reach a steady performance state. Their performance on the discrimination tasks was set to 75% manipulating the contrast of the stimuli, using a staircase procedure. For each trial, the target contrast increases whenever an incorrect response was made, and decreases after two or three consecutive correct responses. The staircase adaptations were larger for the initial practice trials (step size at 25%), followed by a constant step size of 5%, used in the experiment blocks. Participants were instructed that the location of the figures was randomised. After ensuring a fair steady performance of 75% accuracy with as sufficient practice blocks as needed, cTBS was administered and the real experiment began immediately after. After cTBS administration, participants were required to perform three blocks of real/illusory tasks followed by three more of the other tasks. Each block comprises 100 trials. A short break of one minute follows at the end of each block. The order of the tasks and brain sites was counterbalanced across participants and sessions (i.e. cTBS sites). This study was pre-registered prior the research being conducted (https://osf.io/78b3x/).

### Statistical analyses

In order to determine whether cTBS negatively impacts metacognitive awareness, directional (i.e. one-side t test) and planned comparisons are implemented for both overall visibility ratings and metacognitive efficiency values and taking S1-cTBS as a control condition. These planned and directional comparisons always assess whether S1-cTBS visibility ratings or metacognitive efficiencies are higher than DLPFC or PPC-cTBS. All statistical follow a frequentist repeated measures ANOVA followed by a Bayesian implementation calculating the Bayes Factor (BF_10_) representing the relative strength of evidence for the alternative with respect to the null hypothesis. For other statistical analyses (such as accuracy or response bias) no particular directionality is assessed and thus all statistical analyses are ad-hoc with Bonferroni corrections on Type1 errors. All statistical analyses were corrected if parametric assumptions of centrality or sphericity were violated with the corresponding non-parametric alternative or Greenhouse-Geisser correction correspondingly. All data analysis scripts were pre-register prior the research being conducted on the OSF repository (https://osf.io/78b3x/).

## Results

The overall levels of accuracy were not statistically different neither between real and illusory figure tasks (*F*(1, 20) = 0.55, *p* = 0.47, BF_10_ = 0.42), nor between cTBS administration sites (*F*(2, 40) = 0.09, *p* = 0.91, BF_10_ = 0.14; see Figure 3A). The mean contrast level to keep accuracy at 75% did not change across cTBS sites (*F*(2, 40) = 0.01, *p* = 0.99, BF_10_ = 0.13), nor between illusory and real figure tasks (*F*(1, 20) = 0.99, *p* = 0.33, BF_10_ = 0.47). Similarly, reaction times were not observed to differ across cTBS sites (*F*(2, 40) = 0.22, *p* = 0.80, BF_10_ = 0.15), between task types (*F*(1, 20) = 0.06, *p* = 0.81, BF_10_ = 0.30), but were, nonetheless, faster across blocks (*F*(1.45, 29.08) = 13.07, *p* < 0.001, 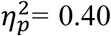, BF_10_ = 0.04), presenting with a linear decrease in the polynomial contrast (*t*(20) = 4.86, *p* < 0.001, *Cohen’s d* = 0.24, BF_10_ = 34.54). Discrimination task sensitivities (d’), with a mean of 1.43 (SD = 0.18), were not affected by cTBS administration (*F*(2, 40) = 0.05, *p* = 0.95, BF_10_ = 0.14) and did not differ between illusory and real figure tasks (*F*(1, 20) = 1.89, *p* = 0.18, BF_10_ = 0.71; see Figure 3B). Similar negative results were also found for the response-free criterion (meta-ca; *F*(2, 40) = 0.80, *p* = 0.46, BF_10_ = 0.23, and *F*(1, 20) < 0.01, *p* = 0.97, BF_10_ = 0.31 respectively), with a mean of -0.02 (SD = 0.17).

**Figure 3.**
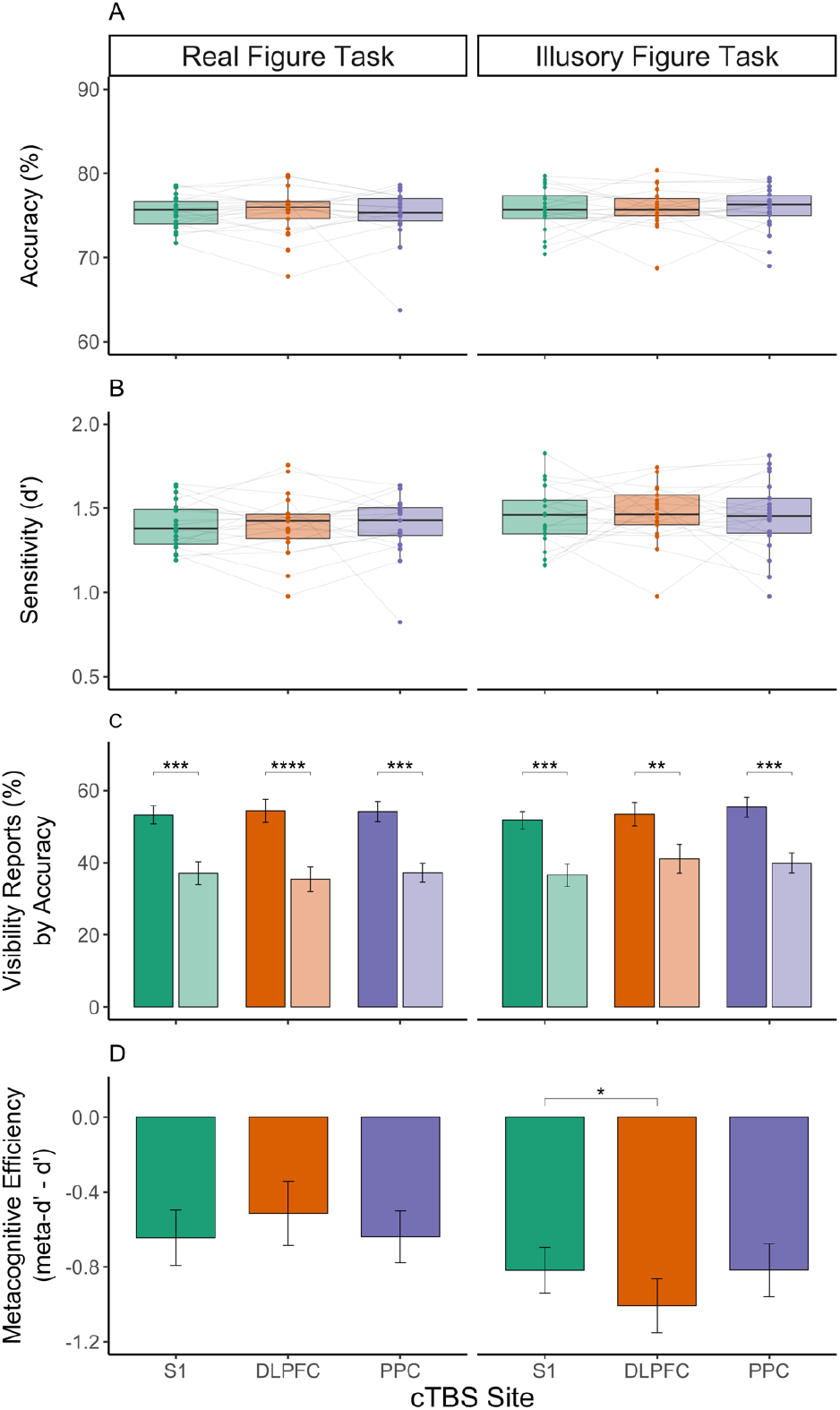
(A) Discrimination task accuracy was successfully stabilised with the use of an adaptive staircase. (B) Which also stabilised perceptual sensitivity (d’). (C) Visibility reports, in turn, were allowed to vary and were observed to be particularly higher at correct (plane bars) compared to incorrect trials (opaque bars). (D) Interestingly, cTBS administration significantly reduced metacognitive abilities when applied to the DLPFC only at the illusory figure task. * = *p* < 0.05, ** = *p* < 0.01, *** = *p* < 0.001, **** = *p* < 0.0001

Metacognitive ratings of visibility were significantly lower for incorrect trials (*t*(20) = 5.44, *p* < 0.001, *Cohen’s d* = 1.19, BF_10_ > 99; see Figure 3C), without differences observed between illusory and real figure tasks (*F*(1, 20) = 0.05, *p* = 0.83, BF_10_ = 0.31). The mean phi correlations between discrimination accuracy and visibility ratings did not change neither between cTBS sites (*F*(2, 40) = 0.05, *p* = 0.19, BF_10_ = 0.13), nor between tasks (*F*(1, 20) = 1.87, *p* = 0.19, BF_10_ = 0.60), with an average of 0.14 (SD = 0.11). The overall ratings of visibility did not change neither due to the nature of the task being performed (*F*(1, 20) = 0.05, *p* = 0.83, BF_10_ = 0.31), nor due to the cTBS administration site (*F*(2, 40) = 0.77, *p* = 0.47, BF_10_ = 0.23). Even though there was an increase in visibility ratings specific to the illusory figure task, the planned contrast on cTBS site (S1 vs DLPFC, PPC) was not significant (*t*(40) = 1.17, *p* = 0.25, BF_10_ = 0.29), concluding that cTBS did not affect reports of visibility.

Metacognitive efficiency values were lower for the illusory figure task in comparison to the real figure task (*F*(1, 20) = 4.86, *p* < 0.05, 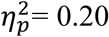, BF_10_ = 1.73), with averages of –0.82 (SD = 0.56) and –0.60 (SD = 0.64) respectively. The order of the first task to perform did not affect metacognitive efficiency results (*F*(1, 20) = 0.05, *p* = 0.82, BF_10_ = 0.31). At the real figure task, no significant effects of cTBS were observed on metacognitive efficiency (*F*(2, 40) = 0.92, *p* = 0.41, BF_10_ = 0.25; see Figure 3D). Planned contrasts revealed that cTBS did not diminish metacognitive efficiency neither when applied to the DLPFC (*t*(20) = 0.99, *p* < 0.84, *Cohen’s d* = 0.22, BF_10_ = 0.13), nor to the PPC (*t*(20) = 0.05, *p* < 0.52, *Cohen’s d* = 0.01, BF_10_ = 0.22). Furthermore, the null effects hypothesis was moderately supported by observed BF_01_ values of 7.99 and 4.57, respectively. At the illusory figure task, the Grubb’s test alerted of a participant presenting a significantly deviant value (G = 2.67, *p* < 0.05), perhaps due to extremely low Type 2 false alarm ratios of 6%, which also affected the normality assumption (W = 0.93, *p* < 0.05). After the removal of this outlier, planned contrasts revealed that cTBS administered over the DLPFC significantly reduced metacognitive efficiencies when compared to the S1 (*t*(19) = 1.89, *p* < 0.05, *Cohen’s d* = 0.42, BF_10_ = 1.96). Administration of cTBS to the PPC did not change metacognitive efficiencies when compared to the S1 (*t*(19) = 0.01, *p* = 0.51, BF_10_ = 0.23). Similar results were observed when obtaining metacognitive efficiency as a ratio (meta-d‘/d’; e.g., cTBS over the DLPFC had negative effects only at the illusory figure task, *t*(19) = 2.15, *p* < 0.05, *Cohen’s d* = 0.48, BF_10_ = 2.95.

## Discussion

This study pursued the objective of investigating whether cTBS over the DLPFC or the PPC could diminish either visibility ratings or metacognitive sensitivities when compared to an active control region (S1) and while sparing objective task performance. Our results indicated that objective task accuracy, discrimination sensitivity and task difficulty were not affected by cTBS administration, neither at real nor at illusory figure tasks. Subjective visibility reports were not affected by cTBS administration site, neither at real nor at illusory figure tasks. The use of illusory contours diminished metacognitive efficiencies irrespectively of cTBS site. Interestingly, cTBS over the DLPFC, not over the PPC, significantly reduced metacognitive efficiencies only while performing the illusory, not the real figure task.

The fact that cTBS administration did not affect visibility ratings supports the idea that, at least for this particular type of perceptual discrimination task using an adaptive staircase procedure to stabilise Type1 performance, visibility reports may not depend on the integrity of the PPN. These null results replicated previous findings with confidence ratings using a similar task design (Bor et al., 2017, 2018). Although a reduction in confidence ratings has been reported when applying online TMS to the DLPFC (Shekhar and Rahnev, 2018), this finding was based on a task without staircase procedure stabilising performance continuously like ours and had a bigger time window between the stimulus presentation and the response time which could have allowed post-perceptual cognitive processes (i.e., a read-out function) to create such effects. It is also important to clarify that our results do not go against the idea of an involvement of the DLPFC or the PPC in other aspects of metacognition such as in mnemonic or in confidence metacognitive tasks.

We attempted to replicate the originally proposed task while introducing improvements that addressed several design concerns raised by Bor et al. (2017). We incorporated the best aspects of both designs into our task. To begin with, this study kept using visibility ratings as originally proposed by Rounis et al. (2010) rather than confidence as introduced by Bor et al. (2017), since we were investigating the subjective sensation of visibility rather than confidence. Second, we implemented practice blocks to ensure that all participants were confident and proficient in their objective discrimination performance by implementing as many practice blocks as they needed, in order to ensure that the observed changes were specific to subjective perceptions of visibility. Third, we continued to use 5% increments in our adaptive staircase to allow for quick adjustment of task difficulty following Rounis et al. (2010). In addition, we have set similar exclusion criteria to those used by Bor et al. (2017) since we agree that unstable datasets may affect the validity of the results. As a result of unstable datasets, three participants were excluded. There were 36% of participants excluded from the included sample, but this was primarily the result of participants not meeting the required accuracy levels of at least 60% correct decisions, which indicated that their visibility ratings would be difficult to interpret with such low Type1 performance levels. As a final point, another significant difference between previous studies was the use of anatomical images to precisely locate the regions of stimulation.

It is important to note that when performing the real figure task initially proposed by Rounis et al. (2010), cTBS administration did not demonstrate any negative effects on metacognition, which replicates previous null findings (Bor et al., 2017; Ruby et al., 2018). This finding does not contradict the notion that prefrontal and parietal regions play a causal role in conscious access and visual metacognition. The lack of effect is consistent with current findings relating visual metacognition to more anterior regions of the prefrontal cortex. However, we raise important doubts regarding the causal role of the PPN in particular, at least when using bilateral cTBS in a theoretically valid metacognitive discrimination task. Many neuroimaging studies indicate that the anterior prefrontal cortex (aPFC) also plays an important role in visual metacognition. Neuroimaging studies have demonstrated that prefrontal cortex gray matter volume and neural activity in healthy participants are positively correlated with confidence ratings and metacognitive sensitivity (Fleming et al., 2010, 2012). Additionally, noninvasive stimulation studies have demonstrated enhanced meta-ratios in perceptual metacognition after online aPFC TMS (Rahnev et al., 2016; Shekhar and Rahnev, 2018), as well as increased memory awareness after aPFC TBS rather than DLPFC stimulation (Ryals et al., 2016). Thus, we cannot exclude the possibility that the aPFC plays a crucial role in metacognition while the DLPFC serves as a supporting component.

Even though all measures of objective task performance were equal between the two tasks, metacognitive efficiencies were lower at the illusory figure task. To our knowledge, this is the first time that Kanizsa illusory contours have been matched to a real figure task and showed an effect on metacognition. It has been argued that illusory figures may crucially depend on top-down priors generated at higher cortical regions, such as the lateral occipital cortex (Murray & Herrmann, 2013). We hypothesised that this effect is due to a reduction in the bottom-up reliability of illusory stimuli which goes in line with previous findings on the role that expectations have over metacognition (Sherman et al., 2015). However, more work is needed in order to confirm that illusory percepts are in fact less reliable than other figures.

Crucially, when performing the Kanizsa illusory figure task, a significantly negative impact of cTBS over the DLPFC, but not over the PPC, was observed for metacognitive efficiency which replicates the original effect reported by Rounis et al. (2010) and would go in line with the idea that the prefrontal cortex is involved in metacognition (Brown et al., 2019; Rahnev, 2017). Despite we found that the DLPFC cTBS selectively interfered with metacognitive efficiency in the illusory figure task, we cannot exclude the possibility that the DLPFC is not directly related to metacognition (Shekhar and Rahnev, 2018), but rather to other functions required for this particular task, namely attention (Buschman & Miller, 2009; Brandt et al., 1998; Corbetta & Shulman, 2002) and the construction of contextual expectations or priors from a top-down perspective (Rahnev et al., 2011; Banica & Schwarzkopf, 2016; Kok et al., 2016). It is important to note that this study used a Kanizsa figure task because of its similarity to the original task, which would help validate systematic effects, however further investigating the influence of the DLPFC guided priors/expectations on metacognitive awareness is required in order to validate our hypothesis that the DLPFC is directly linked with perceptual priors and only indirectly linked to metacognitive awareness.

There are several limitations in the current replicational study. From a behavioural perspective, an important difference between our study and that of Rounis et al. (2010) is that our overall metacognitive efficiencies were low, with M_ratio_ (meta-d‘/d’) lower than 70%. This meant that a significant amount of sensory evidence available for the visibility rating was already lost when making the decision (Fleming and Lau, 2014), leaving little room for cTBS to further dampen it. We understand that the discrimination task was difficult to perform and participants might have preferred to focus on responding accurately, thus downplaying the importance of the subjective feeling of visibility. Similarly, even though we followed the original instructions, the term “visibility” is somewhat a vague concept leaving open the room for alternative interpretations. Participants could have focused on the relevant features of the task at hand, such as seeing clearly where the diamond and square are, or participants could have focused on how generally detectable stimuli are in the task. However, the fact that a significant effect of cTBS was found at the illusory figure task is intriguing because the overall metacognitive efficiencies were lower than those of the real figure task. Rather than isolated, low metacognitive efficiencies have also been reported able to reveal cTBS effects (Lapate et al., 2020). This could potentially reveal that the differences between the two tasks could be explained by the illusory figure task being either more difficult to perform, which would be difficult to argue since the overall discrimination performance was matched with use of the staircase, or as needing extra cognitive resources.

It could still be argued that the DLPFC cTBS effect observed for the illusory figure task may reflect random chance, due to insufficient sample size and statistical power. The idea that cTBS cannot disrupt metacognitive awareness when applied to the DLPFC or the PPC strikes us as the more plausible alternative, especially because we observed a Bayes Factor that moderately supports the null hypothesis for the real task and because Bor et al. (2017) also registered null findings. In this respect our findings are consistent with Bor et al. (2017). Of course it is important to acknowledge that the present study is less than optimally powered, so the possibility of a falsely obtained null result cannot be ruled out (Ruby et al., 2018). Further research is necessary to validate our results.

From a technical perspective, the implementation of cTBS may also have introduced a number of limitations to this study. A major difference from Rounis et al. (2010) is the use of a sham control instead of a one-week apart active control region. The original design can be used to reduce some variances between sessions (mainly pre- and post-cTBS trials), however practice and fatigue can result in additional limitations when comparing visibility reports, as well as a lack of accounting for tactile sensations by using sham stimulation. We attempted to control for these variables by including an active control site, S1, and by using practice trials at the beginning of each session, but the larger between-session variances could potentially be introduced in the current study. Another important point to address is that of the coordinates selected for this study since both the DLPFC and the PPC are big regions encompassing subregions. Rounis et al. (2010) and Bor et al. (2017) defined the DLPFC and the PPC targets as being 5cm anterior or posterior to the “motor hot-spot”. Instead, we followed previous anatomical findings in order to precisely locate the regions of the DLPFC previously linked to metacognitive awareness and functionally connected to the PPC. The following issue is about TMS intensity. An important difference between Rounis et al. (2010) cTBS administration and ours is that we use an intensity adjustment method based on the distance between scalp and cortical surface (Stokes et al., 2005, 2007). As a result of the anatomical characteristics of our sample, the DLPFC was located much closer to the scalp than either the S1 or the PPC. Consequently, the cTBS intensities administered to the DLPFC were approximately half those administered to the posterior regions. Null findings could be attributed to technical issues due to such low intensities. However, it is important to note that the real and illusory figure tasks were performed temporally closely, one after the other, immediately after cTBS was applied to the DLPFC. While balancing the order of the tasks for each participant, the presence of effects at the illusory figure task, but not at the real figure task, suggests the correction method may not have strongly influenced the results.

In conclusion, this study adds new evidence to the ongoing debate regarding the causal role of the DLPFC and the PPC in metacognitive awareness (Rounis et al, 2010; Bor et al., 2017; Boly et al., 2017; Odegaard et al., 2017). Our substantial observed null results in the real figure task go in line with those of Bor et al (2017). Neither the DLPFC nor the PPC cTBS affected neither metacognitive reports nor ability in the real figure task. We thus raise important doubts on the hypothesised direct causal role that the PPN in particular, but not other parts of the frontal cortex such as the aPFC, may play in subjective reports, at least when stabilising objective performance. Nevertheless, we found that the DLPFC cTBS selectively interfered with metacognitive efficiency in the illusory figure task. We hypothesise that when the task involves forming an illusory contour with limited external information, it is necessary to rely heavily on the information generated by internal models (Parr et al., 2018; Yu, 2014). Interestingly, metacognitive ability is a process that requires internal models to generate an evaluation of an individual’s objective performance without external feedback being available. Although the relationship between the two internal models generated in the context of the illusory figure and metacognitive ability remains unclear, limited external information from perceptual and metacognitive processes was found to amplify the impact of DLPFC cTBS on metacognitive efficiency. This result fits with previous studies addressing the DLPFC as a seat of expectation and model creation and thus further evinces that the PPN may not be causally linked to conscious access, at least with use of metacognitive reports. However, legitimate concerns related to less than optimal statistical power, certain aspects of the task and cTBS administration designs, and data quality (such as low metacognitive efficiencies) imply that rendering strong conclusions is inadvisable at this stage. Further research would be necessary to confirm the validity of our findings.

## Acknowledgments

Special thanks to Dr. Daniel Bor and Prof. Hakwan Lau for sharing their Matlab scripts for the real figure discrimination task, as well as to Dr. Niall W. Duncan for building the scalp-distance tool to allow us to estimate the distance from stimulation points on the scalp to the cortex (https://github.com/nw-duncan/scalp-distance). We would also like to thank Ko-Ping Chou, Hsin-Yi Wang, and Paul Z. Cheng for their assistance in collecting data.

This work is sponsored by grants from the National Science and Technology Council, Taiwan (109-2410-H-038-010; 111-2410-H-038-009-MY2) to TYH; (111-2410-H-038-012) to TJL. This work was also supported by the Taiwan Ministry of Education Higher Education Sprout Project.

## Author contributions and competing interests’ declaration

AM and TYH designed the study. AM coded all the scripts, collected and analysed the data. AM, TY and TL wrote, revised and approved the submitted manuscript. The authors declare no conflicts of interest.

## Supplementary information

### S1: Calculation of the adjusted Active Motor Threshold (adjAMT)

The motor evoked potential (MEP) was elicited by placing the coil oriented 45 degrees to the coronal plane and measured from the right first dorsal interosseous (FDI) hand muscle using electromyography (MP160, BIOPAC). The AMT was defined as the lowest stimulator output in percentage that elicits 5 out of 10 twitches of more than 200 μV peak-to-peak amplitude in the contralateral hand, while the participant maintains 20% of a finger-thumb contraction. Previous studies (Stokes et al., 2005, 2007) have shown a positive relationship between scalp-cortex distance and motor threshold. Thus, the AMT found by the previous procedure was adjusted for differences in scalp–cortex distance between brain regions according to a simple linear formula:

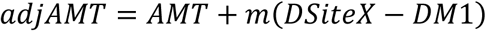

Being the adjAMT the adjusted AMT, AMT the unadjusted AMT, DM1 the distance between the scalp and the motor cortex (M1), and DSiteX the distance between the scalp and the target brain region, either DLPFC, PPC or S1 in this study. Finally, the m is the average distance gradient relating AMT to distance which we defined as 3 based on Stokes et al. (2005). Distances between stimulation points on the scalp and the individual’s cortex were calculated using the *scalp-distance* tool (https://github.com/nw-duncan/scalp-distance).

### S2: Data inclusion

As determined by the standard safety protocols used for MRI and TMS experiments, participants were included if they were 20 years of age, not pregnant, not suffering claustrophobia, had no back problems that would interfere with their ability to recline in the scanner, had no ferromagnetic metallic or electronic implants in their body, had no medical history of convulsions or syncope, nor a family history of epilepsy, and they were not using any medications that were contraindicated for scanning (Hallett, Rossini & Pascual-Leone, 2009). Participants were recruited with normal or corrected-to-normal visual acuity, and without any neurological or psychiatric history.

### S3: Data exclusion

Participants’ objective performance was adjusted by using adaptive staircase to ensure their accuracy was approximately 75%, at each session of the experiment (Kaernbach, 1991). Accuracy levels below 60% would indicate that, for that block, the participant’s performance is at chance level and would, therefore, need to be excluded from further analyses. If any participants had unstable session data sets (across 3 blocks), meaning type1 or type2 false alarm rate below FA <5% or hit rate above 95%, was excluded from analysis.

Treatment of outlier data: Following a recent cTBS meta-analytic study estimating a rather small effect size over the DLPFC (Lowe et al., 2018; which we approximate to a *Cohen’s d*_*z*_ of 0.38) implying that extreme outlier values might shadow our parametric mean comparisons, we run the Grubb’s test (Maximum Normed Residual test) as a rule to determine if any minimum or maximum value should be regarded as outlier and thus excluded from that particular statistical test.

